# Can ERPs and neural oscillations be used to differentiate skilled and unskilled player performance in an ecologically valid sporting model?

**DOI:** 10.1101/2024.11.09.622780

**Authors:** Mike Winstanley, Damian Cruse, Barry Drust

## Abstract

This study aimed to identify neural correlates of high performance in ecological valid, sporting tasks and quantify the differences between high skilled and low skilled players. To do this, Esports was chosen to be the sporting model to facilitate clean brain activity recordings whilst playing sport. After being separated based on skilled objective classification measure, players brain activity was then recorded as they completed a visuomotor psychophysics task and Esports aim-training tasks. Results indicated that skilled players display enhanced response in time-domain activity of occipital parietal electrodes, displaying increases to p100 and p300 peak amplitude. Peak decoding using MVPA, classifying brain activity between skilled or unskilled players, is achieved during the first 300ms of the response. Furthermore, skilled players show greater modulation to visual attention, gated by alpha oscillations localized to the visual cortex, and to cognitive workload, gated by theta oscillations localized to the frontal-midline. During complex, Esports tasks however, skilled players show an increase in frontal-midline theta showing the increased drive of ACC facilitating repeated execution of precise movements. Due to the nature of the model, applied implications are suggested for both coaches and athletes of all sports to utilize this research.

## Introduction

Elite sport forces humans to seek the pinnacle of physical and cognitive performance, making it an interesting phenomenon of human behaviour. Understanding how brain activity relates to performing high-skill motor tasks required in sport, is fundamental to developing better training and development strategies (Thompson et al., 2008). Due to the nature of many traditional sports, recording from the brain presents many challenges that are not easily overcome (Karlinsky et al., 2017). This is due to several factors such as: amplifier dependence, movement (and saccade) artefacts and physical contact with the electrodes (Rowan, 2003).

To address this a new sporting model is proposed using Esports, a new sport where players compete on videogames but played in a competitive way. At the highest level of competition, Esports players participate in a strikingly similar environment to cognitive electrophysiology research. Being played indoors on a computer/console and in a highly controlled environment, lends the sport to being a fruitful model for isolating neural correlates of performance. Research combining cognitive neuroscience and performance in Esports/videogames has been established for decades and identified several important relationships.

Firstly, Esports athletes and videogame players (VGPs) displayed greater speed and accuracy than novices in target detection paradigms which were associated with a shorter latency and larger amplitude p300 component (Mishra et al., 2011). p300 amplitudes have been found to predict visual perception performance (Eimer and Mazza, 2005; Salti et al., 2012; Rutiku et al., 2015), an effect that is abolished through visual blurring of targets (Heinrich et al., 2010). Source analysis reveals that the increase in p300 amplitude originates in occipital-parietal regions (Bablioni et al. 2006). A variety of sources position p300 as a manifestation of conscious access, not simply visual perception, and contains signaling to report task-relevant stimuli (Pitts et al., 2014; Mashour et al., 2020). In this way, p300 can be seen as a reflection of the demand for attentional resource allocation in response to sensory stimuli (Gray et al., 2004). Amplitude increases in the p2/300 component of parietal networks have been found after video game training (Wu et al., 2012) but show a negative correlation with game difficulty in expert VGPs (Allison and Polich, 2008).

Furthermore during an Esports based visual discrimination tasks, professional Esports players displayed stronger stimulus-locked alpha-band power (Gostilovich et al., 2023). Alpha power shows strong links to visual detection performance through an inverse relationship (Benwell et al., 2017; 2022). Specifically, the ability to discriminate the presence of a visual stimulus decreases with alpha power (Van Dijk et al., 2008). Importantly, low alpha power was also associated with significantly higher ERP amplitudes (Iemi et al., 2017). The associated power decrease in pre stimulus alpha is resulting from enhanced global visual system excitability (Lange et al., 2013).

Finally, increases to frontal theta power during VG play compared to rest periods (Pellouchoud et al., 1999) have been observed. Theta power is associated with activity in the anterior cingulate cortex (ACC) and prefrontal cortex in source localization studies (Onton et al., 2005; Ishii et al., 2014). Theta power localized to the ACC and other frontal structures has a diverse range of functions in complex cognition such as action regulation (Luu and Pedersen, 2004) and monitoring (Cavanagh et al., 2009); conflict monitoring (Botvinick et al., 2004), task selection (Womelsdorf et al., 2010). During Esports competition, theta power increases as a function of rounds progressed and increases prior to the onset of performance feedback (Sheikholeslami et al., 2007). In an older population, VG play has been shown to significantly increase theta power which was correlated to performance improvements on a plethora of cognitive tests (Anguera et al., 2013).

This paper seeks to further our understanding of the relationship between visual processing, cognitive processes and performance in sport. To achieve this, Esports is chosen to function as model for all sport that overcomes well established limitations of recording brain activity during active sport. Two experiments are proposed that directly test the fundamental movement in Esports, fast and reflexive aiming. The first is a psychophysics task that uses simple, high contrast targets and induces discrete aiming movements pulling participants across the screen. The second is performed on a commercial ‘aim-trainer’ (AimLabs^TM^), increases the complexity of the visual information with coloured, 3-dimensional targets and increases the complexity of the aiming movement by inducing sequential movements. The experiments use 64-channel EEG to record brain activity, but the experiments differ in how the activity is processed. In the psychophysics task, custom programming facilitates evoked brain activity however, in the second, only induced activity is available due to the commercial design of the game, specific events beyond the beginning and the end of the task are not able to be marked. Using this data, we examined whether performance on these tasks can be used to differentiate skilled from unskilled players. Secondly, we examined whether brain activity recorded during the performance of psychophysics task, looking specifically at pre-stimulus alpha and post stimulus ERP components (p1/300), can explain the observed performance difference. Finally, we examined whether induced theta power is upregulated during active performance and determine to quantify its relationship to performance. In this way, we build an argument for how skilled players execute a higher level of visuomotor and Esports performance with an enhanced response to visual information, modulations to the task period dependent cognitive processes of attention and workload, and enhancements to movement related neural oscillations.

## Methods

### Participants

For this study, 37 participants (15 Male, 12 Female) took part in the experiment. 36 of these were students at the University of Birmingham and one was a Marine (UK Armed Forces). Of the participants, 19 were classified as Skilled players and 18 were classified as unskilled players based on the methods described above. After reporting Esports playing time based on a self-report questionnaire, populations were determined based on Esports experience only. Players who played video-games, whichever input modality, but not Esports, were determined to be inexperienced. After completing various Esports and psychophysics tasks, their performance on these tasks was used to perform k-means clustering and identify clusters of similarly performing player groups. Silhouette scores determined that two clusters were optimal, and players were classified based on their cluster. Their previous experience was not included as part of the classification process. All participants were right-handed, didn’t wear glasses and had no history of neurological disorders. Crucially, all participants reported some experience with videogames. The experimental procedure was approved by the ethics committee review in the School of Psychology (University of Birmingham) and informed consent was obtained from all participants before participation and checked again after completion. Participants were compensated for their time. It was not appropriate to involve participants in the design, or conduct, or reporting, or dissemination plans of our research

### Experimental Procedure - Psychophysics

**Figure 1.**
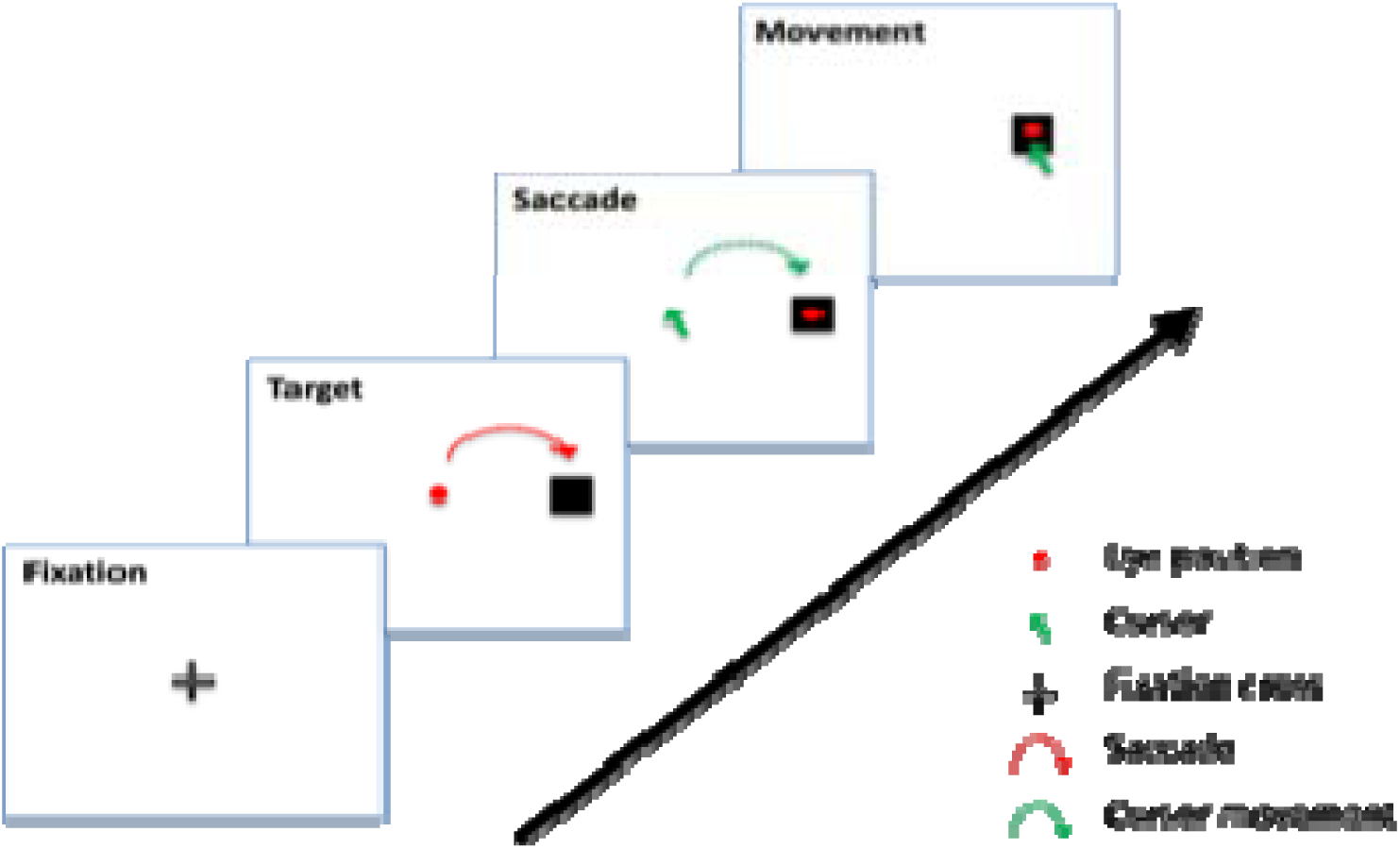
Experimental design for the experiment displaying the fixation period, stimulus onset, eye movement and cursor movement.

The psychophysics experiment was created using SR Research experiment builder to accurately co-register with the eye-tracker and stimulus pc. In each experiment, a fixation cross began the trial by being present for 1000ms localizing eye fixation to the center of the screen. After the fixation period was over, the mouse location was set to the center of the screen and the response target, a visual stimulus, would appear on the screen. The target was a small black square, projected on a white background to maximize contrast. Participants would then have 1000ms after stimulus onset to move the mouse to the target and click on it, with only clicks registering within the target boundary being registered. The reaction time, calculated from stimulus onset to a successful click, was recorded, as well as the number of failed trials, reported as errors. If the trial was successful, a blank screen was present for 500ms. If the trial was failed because a successful click wasn’t registered within the 1000ms this would be recorded as an error and a message reading “FAILURE, MOVE FASTER!” would appear on the screen in bold red lettering. After 500ms of this message being present, another trial would begin, denoted by the fixation cross reappearance. There were three different blocks, with each block using a different target size decreasing by 10 pixels squared each block (30x30 pixels to 10x10 pixels). All targets remained the same colour throughout and occurred at a set number of locations (20 left and 20 right, 10 center).

### Experimental Procedure – Esports Aiming Test

**Figure 2.**
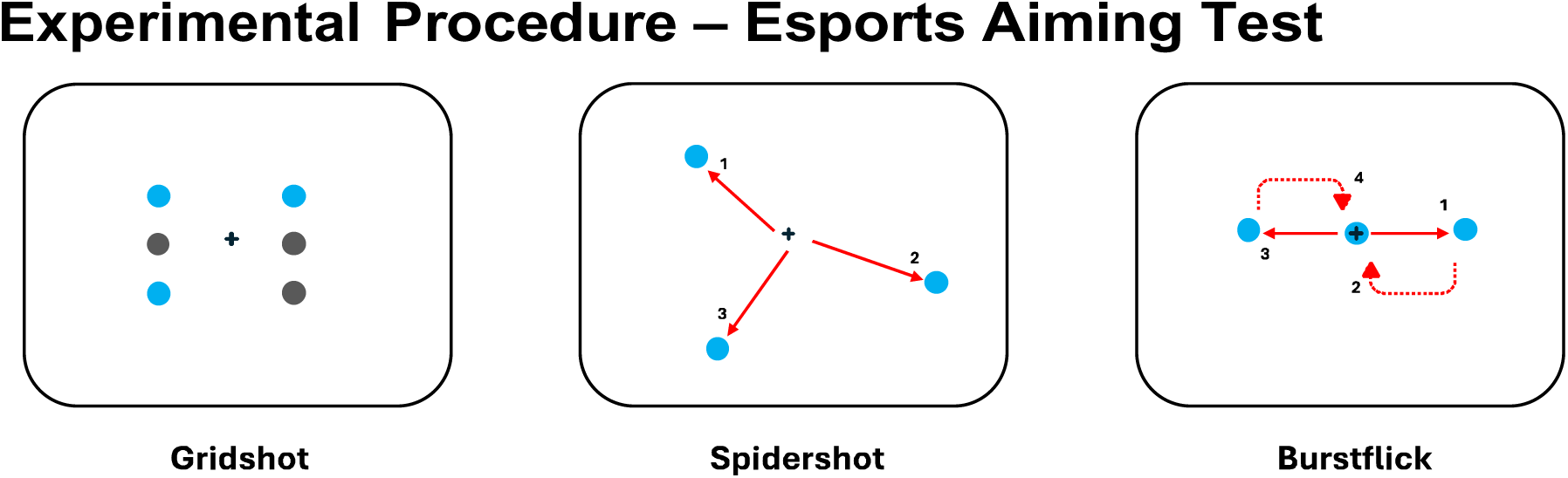
Schematic of each Esports aiming tasks: Gridshot, Spidershot and Burstflick. The blue circles are the targets, grey circles are possible target locations one others are destroyed, the black cross is the crosshair denoting aim position, red arrows denote crosshair movement towards each target, numbers denote the order in which movement were made.

To test performance in an Esports related aiming task, the commercial aim-trainer AimLabs was used. Participants took part in three different tasks which were designed to train performance of aiming movements in Esports. All tasks had slight differences between each other to test gather a more holistic understanding of aiming performance. Fundamentally, each task tested performance in the same way, clicking inside the boundary of the target ‘destroys’ it and counts towards the score defining overall performance in the task. After a target is destroyed, another appears somewhere else on screen subject to the differing task conditions. Each task lasts 60 seconds long and players must generate as high of a score as possible. Gridshot is the first task completed by players. It consists of 3 targets appearing on screen at a time in a 3x2 grid shape. Once a target is destroyed, another appears somewhere else in the grid, continuing until the task is complete. Spidershot is the second task and consists of a singular target appearing randomly on the screen. Once destroyed, another target appears randomly on the screen, continuing in this vein until the task is complete. Finally, Burstflick is the final task. It consists of a target appearing in the center to be destroyed, once this has occurred another target appears for 1 second laterally from the central target. Once this has been destroyed or the timer has run out, another central target appears. The task continues in this vein until complete. In all tasks, clicking inside the boundary for each target in succession adds a multiplier to the score.

### Eye-Tracking

Eye-movements were recorded using an Eyelink 1000 plus (SR Research), with eye-tracking calibration steps performed before all experimental blocks before the delivery of tasks instructions. To quantify eye-movements during the study, SR research’s data analysis tool, Data Viewer was used. This facilitated the identification of saccade, fixations, and trial reports. Due to the experimental coding being in another data package from SR research, all trials ported over contained the necessary trial condition information such as the target size, location, and calibration reports. To identify eye-movements exclusively during the trial period, a reaction time variable could be initialized which marked the moment of stimulus onset, detected by the time the host pc received the stimulus onset trigger (the same trigger received by the EEG amplifier) to the moment it received the outcome trigger, either success or failure. As such, only eye-movement event-locked to the stimulus onset were included in analysis. Saccades below 1 degree were excluded from analysis.

### EEG recording and analysis

The data was acquired using a 64-channel EEG (BioSemi) and processed using MNE – python toolbox. The data was subjected to a number of pre-processing and processing steps. Briefly, noisy sensors were first removed and interpolated by RANSAC algorithm. Then the data was downsampled to 200hz to reduce computation time. After downsampling, events were marked depending on population (skilled or unskilled), event (stimulus onset or response) and outcome (success or failure). From here the data was passed through the AutoReject algorithm to reconstruct and drop extremely noisy epochs before being passed to the ICA algorithm which allowed for artefact detection and removal. Then the data was passed for a final time to AutoReject to reconstruct any remaining noisy epochs and drop any that still didn’t pass the threshold (a lower threshold than the first AutoReject pass). At this point, the now clean data was filtered from 1-30Hz (high and low-pass) and average referenced. Epochs from this position could finally be averaged together to form evoked objects, or grand average ERPs, per population, event, and outcome.

In the Esports task, induced activity was recorded. To identify game durations, a trigger was sent at the beginning of the video recording elements. This trigger was recorded in the message log of Weblink and could then be used to isolate the time in the video when the task began. By subtracting the offset between when the trigger was received by the eye-tracker, from the task start, this could serve as the initial crop point. Each task lasted for 60s, so the crop end time was calculated as the task start time + 60s. Each file was read in individually and cropped according to the trial start and end time.

### Time-domain analysis

To identify the visual stimulus related component of ERPs, occipital parietal electrodes were grouped and plotted as ERPs, event locked to stimulus onset, depending on the outcome. Stimulus onset events that were followed by events of interest were defined using the function ‘define_target_events’ allowing for new events to be created if a target event follows the original event, within a certain time-window. This allowed stimulus onset events during success trials, and stimulus onset during failure trials, to be identified separately and compared.

### Frequency-domain analysis

With clean and pre-processed data, Morlet waveform analysis was applied to frequencies of interest. In this experiment, the frequency ranges used were Theta (4-7Hz), Alpha (8-12Hz), Low Beta (13-21Hz) and High Beta (22-30Hz), calculating individual frequencies in steps of 1Hz. To define the number of cycles, the frequency range was divided by 2 and fast Fourier transform was applied. Power for each frequency was then averaged.

### Statistical analysis

Statistical analysis was conducted differently depending on the type of data, behavioral data and outputted measurements of time/frequency elements, were analyzed using Prism (GraphPad). Examples of this are behavioral performance, eye-tracking, ERP measurements and power estimates. Statical analysis of complex time/frequency elements were analyzed using MNE-Python. To establish a statistical relationship between time/frequency elements, outcome, and population, peak measurements of grand averages and averaged power estimates were made. These measurements were then statistically analyzed using a 2-way ANOVA comparing Population x Outcome, corrected for multiple comparisons using Tukey’s post-hoc test. 1-D cluster-based permutation statistics were calculated for difference waves, testing deviation from zero. An F-value threshold of 6 was used to determine a significant cluster.

### Source localization

To achieve source localization several different techniques were used to estimate both activation and power changes in the source space. To compute all solutions, a forward model was created using the standard template MRI subject fsaverage (FreeSurfer). Using Dynamics Imaging of Coherent Sources (DICSs), a volumetric forward model was created. To do this the template MRI was used to construct boundary element model (BEM) using a three-shell model (brain, inner and outer, skull). To calculate source activations, dynamic statistical parameter mapping (dSPM) was created. This uses the minimum norm or weighted minimum norm inverse operator by normalizing its rows. To calculate event-related source power changes the DICS method was used with the volumetric forward model. Cross-spectral density was calculated for each frequency band using morlet waveform transformations using a baseline covariance matrix (pre-trial) and an active covariance matrix (during trial), in this case, the baseline was set 1.5-1s prior to the pre-stimulus fixation cross period, where a blank screen was present after the termination of the outcome message.

## Results

**Figure 2.**
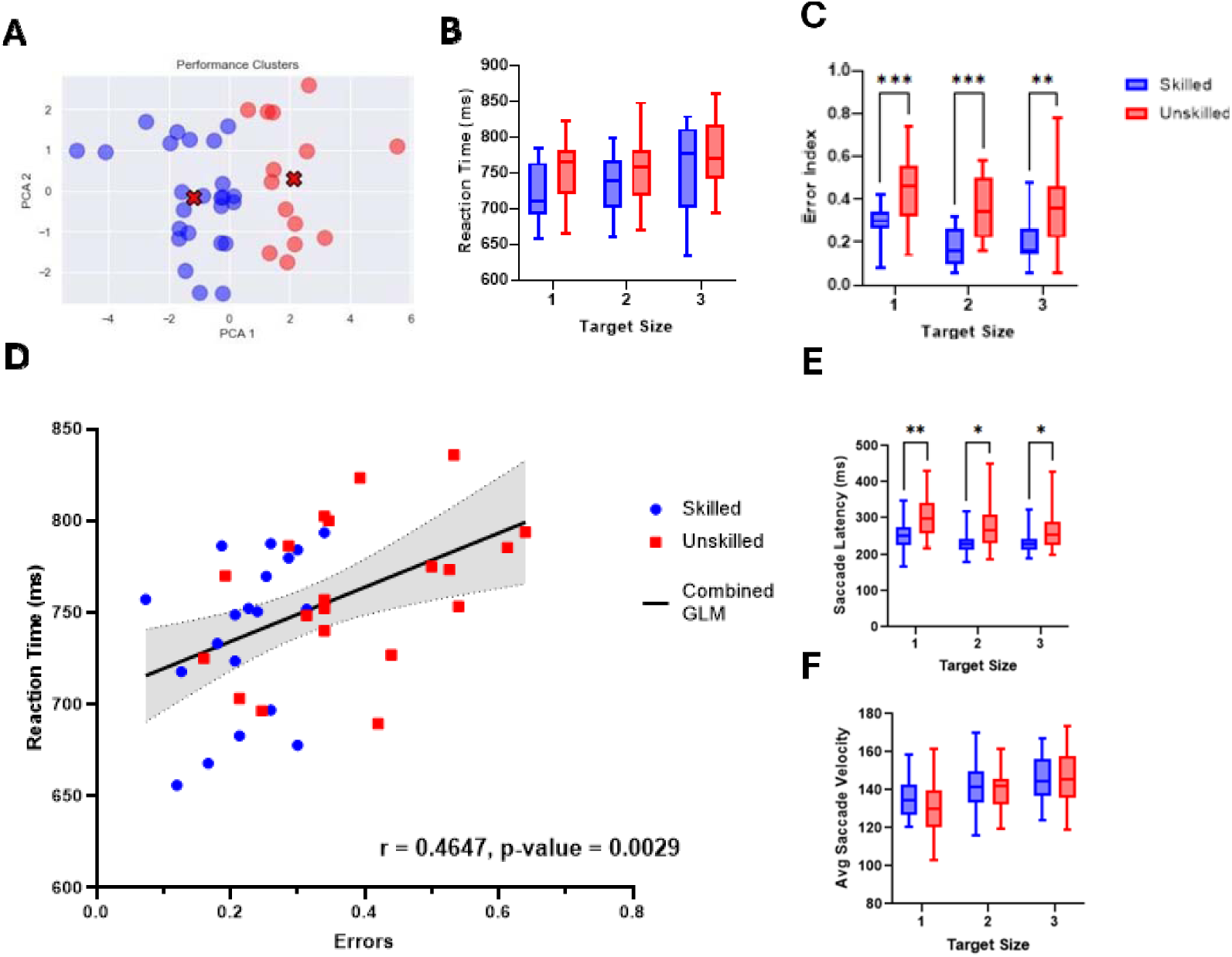
Behavioural and eye-tracking performance across psychophysics tasks. A) The classification output of k-means clustering on Esports performance data, identifying two distinct groups. B) reaction time psychophysics task depending on target sizes (1 = large, 2 = medium, 3=large) with skilled players in blue and unskilled players in red. Upper and lower range are shown and box size represents standard deviation. C) Box plot of reaction time depending on target size. B) Box plot showing error index depending on target size. D) Correlation between reaction time and error index with general linear model plotted in black and error bars coloured in grey. E) Box plots displaying the time to first saccade across populations during the psychophysics task. F) The average saccade velocity of saccades across populations during the psychophysics task. Statistical analysis in box plots is 2-way ANOVA corrected for multiple comparisons using Tukey with significant comparisons marked (* = 0.01, ** = 0.001, *** = 0.0001).

The behavioural results indicated that there is not a significant interaction between target size and population with reaction time (F(2,72) = 0.3235, p = 0.7247), however there is a significant interaction between target size and population with error index (F(2, 72) = 9.457, p = 0.0259). Skilled players displayed faster average reaction times (F (2, 36) = 19.76, p=0.0911) in relation to unskilled players, although the lack of significant interaction does not justify post-hoc comparisons. The number of failed trials, referred to as errors, was used to create an error index. The likelihood of a player to make an error, was significantly different in skilled players compared to unskilled players (F (2, 36) = 19.76, p = <0.0001). Tukey’s multiple comparison post-hoc test indicated significant differences at all target sizes, from 1 to 3 (p = 0.0005, p = 0.0002, p = 0.0016, respectively). Skilled players made significantly fewer errors and thus had a significantly small error index than unskilled players across all target sizes. In the correlation analysis there is a significant correlation between the two performance variables across the combined population (p-value = 0.0029). There is a strong positive relationship between reaction time and error index (r = 0.4647, p = 0.0029). Skilled players showed a significantly better behavioural performance by demonstrating a faster reaction time and fewer errors, two variables that are strongly correlated. Eye-tracking results indicate a difference in execution between skilled and unskilled players (EP and NEP). There was a significant interaction between Population and Target size in the Saccade latency (F(5, 216) = 3.045, p = 0.0112. There was also significance in population difference (F(1, 216) = 15.24, p = 0.0001) and a significant difference across all target sizes (F(5, 216) = 88.31, p < 0.0001). When correcting for multiple comparisons there are significant differences in saccade latencies across with p-values of 0.0032, 0.0059, 0.0394 for sizes 1, 2, 3 respectively. There is not a significant interaction between population and Target size in average saccade velocity (F(5, 216) = 1.824, p = 0.1094). There is not a significant difference across populations (F(1, 216) = 0.2281, p = 0.6334) but there is a significant difference across outcomes/target size (F(5, 216) = 23.47, p<0.0001).

**Figure 3.**
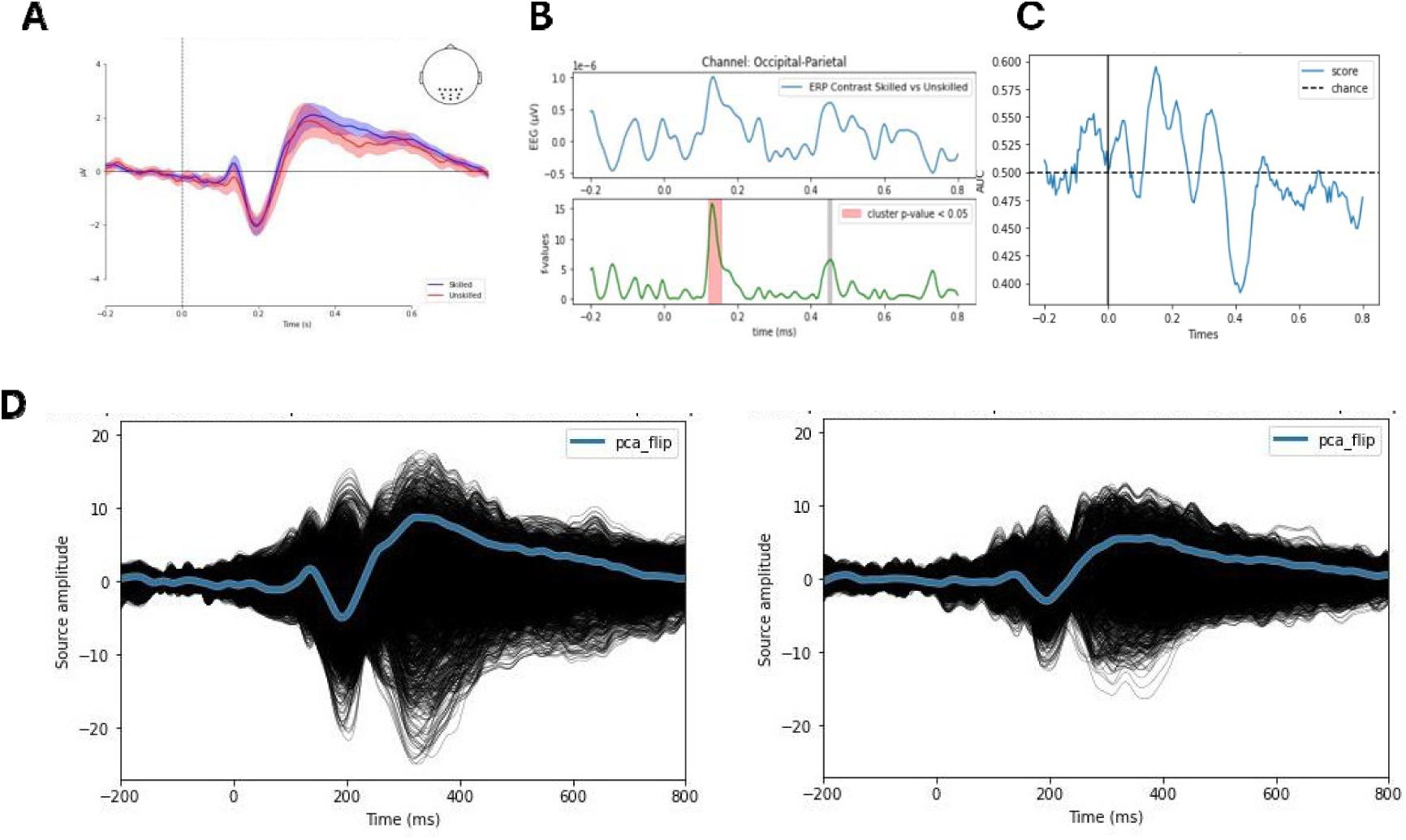
Time domain activity during the stimulus onset period in both the sensor and source space. A) Sensor space ERP from occipital parietal electrodes, skilled players coloured in blue and unskilled players in red. B) Grand average difference wave ERPs between skilled players, unskilled players and between outcomes across populations, corrected for multiple comparisons using cluster permutation test. C) Multi-variate pattern analysis differentiating between skilled and unskilled players during stimulus onset period. Decoding performance is plotted in blue with above 0.5 being above statistical chance of occurring. D) dSPM source amplitude differences between skilled (left) and unskilled players (right) computed with a sign flip across visual cortex labels to avoid signal cancellation when averaging signed values.

Grand average ERP from occipital parietal electrodes shows a number of similarities at the sensor space level. The major difference comes around 100-200ms after stimulus onset with a much stronger p100 present in skilled players. The difference wave plots and subsequent 1-D cluster permutation statistics, display a significant difference period in skilled vs unskilled players between 100-150ms after stimulus onset in successful trials, but no significant difference periods afterwards, although a post threshold, sub significant, peak occurs at 400-450ms. In the failure trials, an above-threshold, but insignificant peak occurs between 100-150ms after stimulus onset, but no significant difference is detected comparing skilled and unskilled players. The source amplitude changes are plotted to transform sensor space voltage changes to the source space. Within both populations, differences in source amplitude are present between outcome conditions, with a greater amplitude occurring in successful trials occurring 300ms after stimulus onset. The peak source amplitude is higher in skilled players compared to unskilled players; however no detectable differences occur in the failure outcome between populations.

**Figure 4.**
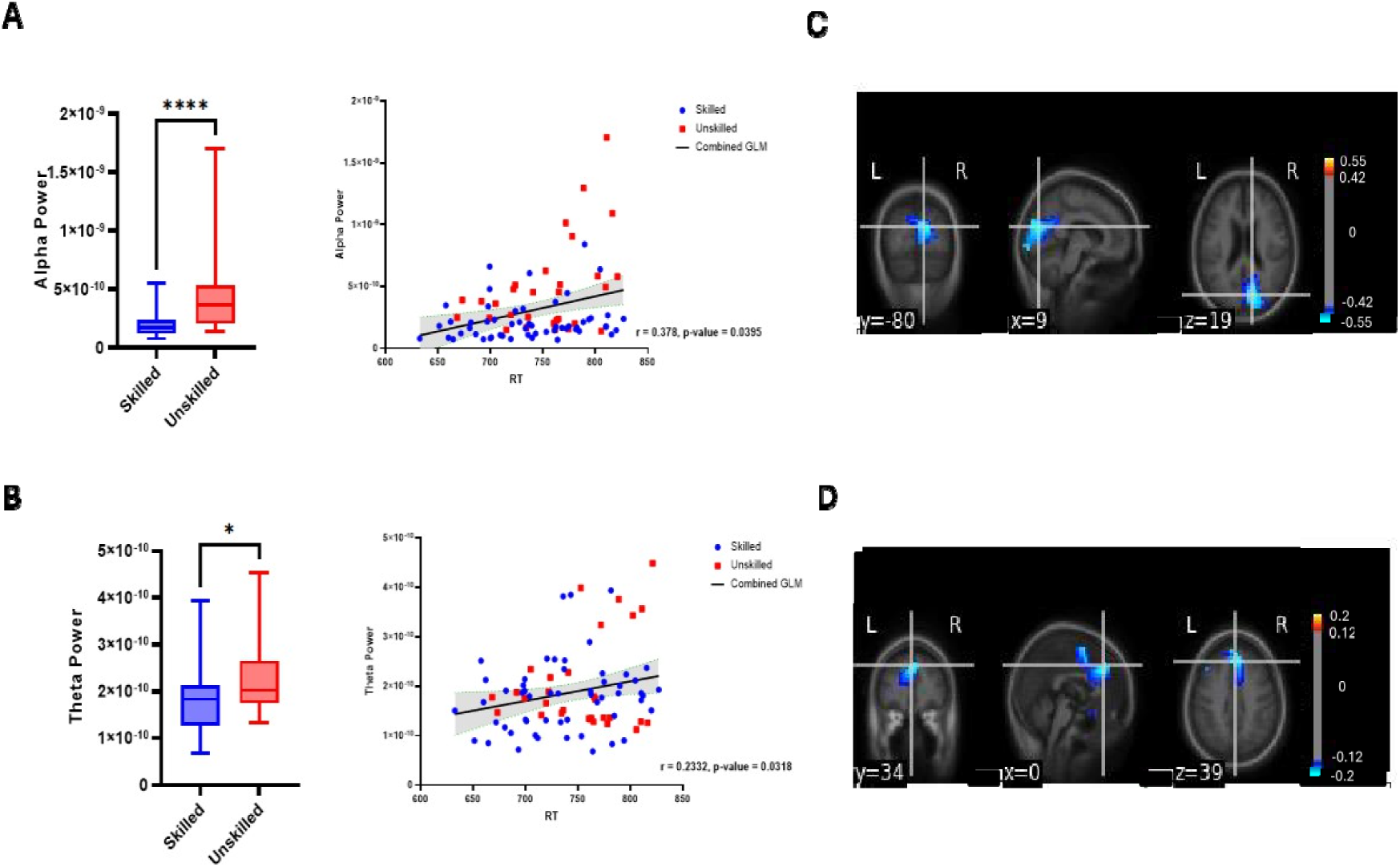
Neural oscillations during pre-stimulus (-0.5 – 0s) and post-stimulus (0 – 0.5s) phases during psychophysics experiment. A) Alpha power differences in the pre-stimulus phase displaying the raw power difference between skilled and unskilled players and the correlation between theta power and reaction time. B) Theta power differences in the post-stimulus phase displaying the raw power difference between skilled and unskilled players and the correlation between theta power and reaction time. C) DICS source localization of alpha power during the pre-stimulus phase projected onto a template MRI showing the difference between skilled and unskilled players. D) DICS source localization of theta power projected onto a template MRI showing the difference between skilled and unskilled players. Statistical analysis in box plots used unpaired, non-parametric t-tests (Mann-Whitney) after failing normality tests (Kolmogorov-Smirnov test). Significant comparisons marked (* = 0.01, ** = 0.001, *** = 0.0001, **** = <0.0001).

To compare neural oscillations during execution of movements, pre-stimulus alpha power and post-stimulus theta power were quantified at timepoints where they impart the most influence. Pre-stimulus alpha power is quantified in a box plot displaying how skilled players show significantly lower alpha power than unskilled players, analysed through unpaired non-parametric t-tests, Mann-Whitney (p= <0.0001). post-stimulus theta power also showed a significant reduction in skilled players, analysed through unpaired non-parametric t-tests, Mann-Whitney (p= 0.0156). Correlational analysis indicated a significant relationship between pre-stimulus alpha power and reaction time (r = 0.378, p = 0.0395) across all participants and in post-stimulus theta power (r =0.2332, p = 0.0318). This indicates a strong inverse relationship between pre-stimulus alpha power and post-stimulus theta power with high performance. There is a strong relationship between neural oscillations and performance, displayed by significantly reduced pre-stimulus alpha and significantly reduced post-stimulus theta.

**Table 1.** Summary table from 3-way ANOVA of theta power comparing task x experimental condition x population.

| ANOVA table | SS | DF | F (DFn, DFd) | P value |
| --- | --- | --- | --- | --- |
| Task | 1.674e-021 | 2 | F (2.000, 64.00) = 1.132 | P=0.3289 |
| Active vs Passive | 2.521e-019 | 1 | F (0.9344, 29.90) = 39.30 | P<0.0001 |
| Skilled vs Unskilled | 1.005e-019 | 1 | F (1, 32) = 4.699 | P=0.0377 |
| Task x Active vs Passive | 6.155e-021 | 2 | F (1.740, 55.67) = 4.732 | P=0.0162 |
| Task x Skilled vs Unskilled | 1.256e-021 | 2 | F (2, 64) = 0.8487 | P=0.4327 |
| Active vs Passive x Skilled vs Unskilled | 4.063e-020 | 1 | F (1, 32) = 6.334 | P=0.0171 |
| Task x Active vs Passive x Skilled vs Unskilled | 3.092e-021 | 2 | F (2, 64) = 2.377 | P=0.1010 |

**Table 2.** Summary table from 3-way ANOVA of theta power comparing task x experimental condition x population.

| ANOVA table | SS | DF | F (DFn, DFd) | P value |
| --- | --- | --- | --- | --- |
| Task | 1.407e-021 | 2 | F (2.000, 64.00) = 1.242 | P=0.2957 |
| Active Passive | 3.168e-020 | 1 | F (0.9266, 29.65) = 5.522 | P=0.0278 |
| Skilled vs Unskilled | 2.613e-020 | 1 | F (1, 32) = 1.365 | P=0.2514 |
| Task x Active Passive | 8.617e-022 | 2 | F (1.964, 62.85) = 0.7502 | P=0.4742 |
| Task x Skilled vs Unskilled | 8.353e-022 | 2 | F (2, 64) = 0.7374 | P=0.4824 |
| Active Passive x Skilled vs Unskilled | 9.350e-021 | 1 | F (1, 32) = 1.630 | P=0.2109 |
| Task x Active Passive x Skilled<br>vs Unskilled | 4.721e-022 | 2 | F (2, 64) = 0.4110 | P=0.6647 |

Across the multiple tasks testing two different fundamental movements in Esports, flicking and tracking, skilled players performed significantly better than unskilled players. In flicking tasks, the interaction between task and population is significant for Score (F (2, 64) = 3.556, p = 0.0343). There was a significant score difference across each task (F (1.059, 42.34) = 109.0, p = <0.0001) and across population (F (1, 40) = 16.55, p = 0.0002). Tukey’s multiple comparison post-hoc test indicated there was a significantly higher score in skilled players across each task (Burstflick, p = <0.0115; Gridshot, p = 0.0001; Spidershot, p = 0.0001). Time to Kill also showed a significant interaction between task and population (F (2, 64) = 3.382, p = 0.0467). Differences across tasks (F (1.788, 57.21) = 58.57, p = <0.0001) and across populations (F (1, 32) = 11.16, p = 0.0021) were also significant. Tukey’s multiple comparison post-hoc test also indicated that there were significant differences in Time to Kill across all tasks (Burstflick, p = <0.0499; Gridshot, p = 0.0023; Spidershot, p = 0.0306). Frontal central theta power increased significantly in active conditions compared to passive conditions (F (0.9344, 29.90) = 36.87, p = <0.0001) but was not significantly different between participants (F (1, 32) = 3.373, p = 0.0756). By comparing both active and passive conditions and skilled and unskilled groups, a significant difference is revealed (F (1, 32) = 4.604, p = 0.0396).Occipital parietal alpha power significantly differed between active and passive conditions (F (0.9266, 29.65) = 5.522, p = 0.0278) but didn’t significantly differ between populations (F (1, 32) = 1.365, p =0.2514). There was not a significant difference by comparing both active and passive conditions and skilled and unskilled groups (F (1, 32) = 1.630, p = 0.2109). In flicking tasks, there are significant differences between theta power levels across populations (F (1, 32) = 4.745, p = 0.0369) but not between tasks (F (1.901, 60.83) = 0.5502, p = 0.5709). Tukey’s multiple comparison post-hoc test indicated significantly higher theta power in two tasks (Burstflick, p = 0.0086 and Gridshot, p = 0.035) but not in Spidershot (p = 0.0919). Occipital-parietal alpha power showed significant differences across population (F (1, 32) = 4.391, p = 0.0441) but not across task (F (1.845, 59.04) = 0.7792, p = 0.4540). Tukey’s multiple comparison post-hoc test indicated significant increases in alpha power in skilled players in two tasks (Gridshot, p = 0.0316 and Spidershot, p = 0.0360) but not in Burstflick (p = 0.0536) although it was statistically trending. Whilst playing an Esports related aim-training task, frontal-central theta power increased significantly in the active conditions compared to the passive condition across both populations. Theta power also increased significantly in skilled players compared to unskilled players. The overall alpha power increased in active conditions compared to inactive conditions, although this effect was not significant and only present in skilled players. There are differences in the oscillatory power exhibited by skilled and unskilled players during the tracking experiments. Skilled players displayed significantly higher theta power across all tasks compared to unskilled players. Unskilled players showed higher alpha power in the active tasks, but the result is not significant.

**Figure 6.**
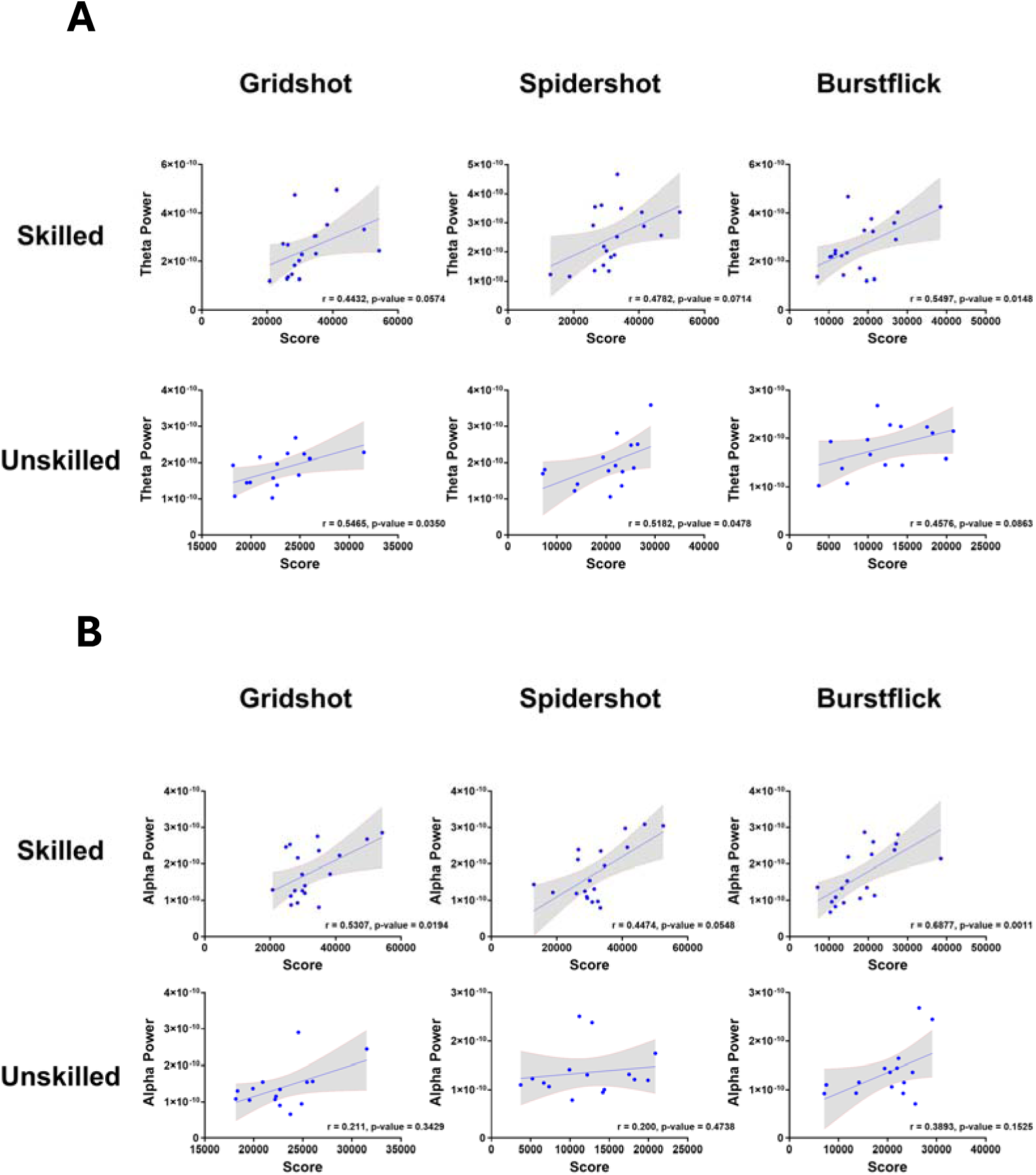
Correlational analysis between neural oscillations and behavioural performance in Esports aiming tasks. A) Correlations between theta power and score across each task in frontal-central sensors in flicking tasks with general linear model plotted in black and error bars coloured in grey. B) Correlations between alpha power and score across each task in frontal-central sensors in flicking tasks with general linear model plotted in black and error bars coloured in grey. R values p values for each correlational are reported.

Correlational analysis revealed significant positive relationship between behavioural performance within each task and theta power. In skilled players this relationship was significant for the tasks: Gridshot, Spidershot and Burstflick. In unskilled players this relationship was strongly positive for Gridshot and Spidershot, but negative for Burstflick with no relationships being significant. In all cases, this slope of the GLM was drastically steeper in skilled players than unskilled players highlighting a much stronger relationship. In frontal central sensors, there was a significantly stronger relationship between theta power and performance, presenting theta power as a indicator of performance in skilled players but not unskilled players. However, by using Fisher’s r-to-z approach, there were no significant differences across all correlations (Gridshot: z= 0.359, p = 0.36; Spidershot z = -0.139, p = 0.445; Burstflick: z = 0.324, p = 0.373). Alpha power was shown to be correlated strongly to performance in skilled players but not in unskilled players. In skilled players, a significant positive correlation was found in the Gridshot and Burstflick tasks and a strong positive correlation was found in Spidershot. In unskilled players, the relationship between alpha power and performance was much weaker, shown by insignificant p-values and smaller correlation coefficients. The strongest relationship was with the Burstflick task, with Gridshot and Spidershot being much weaker (shown by a smaller r value). None of these relationships are significant. Across all players, alpha power seems somewhat correlated to performance, with a stronger correlation in skilled players. However, by using Fisher’s r-to-z approach, there were no significant differences across all skilled vs unskilled correlations (Gridshot: z = 0.612, p = 0.27; Spidershot z = -0.087, p = 0.465; Burstflick: z = 1.133, p = 0.129).

## Discussion

The present study aimed to establish whether brain activity can be used to differentiate players based on their performance within a sporting model, identifying neural correlates that are driving the higher performance level. It has introduced the use of Esports as a sporting model where unifying principles of how information is extracted and utilized to execute a price movement response can be extrapolated to other sports. By applying cognitive electrophysiological techniques to sport science data, logistical challenges in common practices are overcome, whilst retaining ecological sporting validity. To that end, an ecologically valid methodology was employed by capturing participant performance on a fundamental movement psychophysics task and aim-training tasks within the commercial aim-training software, AimLab.

The tasks used induced fundamental movements required by FPS Esports in a high-volume, fast-movement paradigm. In the psychophysics task, discrete movement responses were made to the onset of simple stimuli (black squares and white background) which had to be complete within a short time period. In the aim-training tasks multiple targets were always present on the screen, with participants freely choosing which one to shoot at first, before switching to the next. This increased the complexity of the task drastically and pushed it towards a higher level of ecological validity, more closely emulating scenarios experienced in Esports. Furthermore, the visual information was 3-dimensional, highly coloured and the targets were dynamic. As such, the study encompasses both discrete and sequential movements.

To identify skilled and unskilled players contained within the recorded population, an objective data-driven approach was utilized. In this method, performance data from all experiments was used to test for the presence of clusters in the data creating separate performance groups. Players that cluster together, perform most similarly. In this way, two groups were formed based on performance alone and increased separation when based on experience alone. To address this problem, there needs to be a way of separating based on performance and finding two populations that have a large difference, but also intrinsic show similar performance profiles. A first issue is, however, the number of performance metrics used, and how to choose which one. A solution to this, is to use PCA analysis to project multi-dimension data on common axis, through dimensionality reduction. PCA was applied to all the performance metrics, achieving dimensionality reduction, and re-projecting the data along a 2-dimensional axis of principal components, retaining information and capturing variance derived from the data. To probe whether experienced players perform better, classification was applied to the un-grouped data. Finally, the data groups were re-classified using k-means clustering to attribute the data items, to the optimal two clusters, determined by silhouette analysis. Intrinsically, the data items within these clusters contain participants whose performance is most similar throughout the data set, but also producing maximum separation to the other cluster.

Skilled players performed significantly better than unskilled players across all behavioural metrics across both tasks. Behavioural analysis of psychophysics task performance revealed that skilled players show a reduced reaction time and number of errors compared to unskilled players. These two metrics also significantly correlated with one another, where players with the fastest reaction times displayed the fewest errors. Skilled players also display earlier saccades but with relatively similar average saccade velocity. In the aim-training tasks, skilled players displayed significantly faster Time to kill (TTK) and higher Score, a combination of speed and accuracy, which accumulated throughout the trial resulting from successful shots and amplified by consecutive successes, displayed the strongest significant differences between the populations. Skilled players accumulated a significantly higher score than unskilled players across all tasks.

Neural correlates of this higher level of performance are separated based on skill level across the sensor and source space, emerging from two different domains of brain activity. The time domain, how activity changes over time, and the frequency domain, how activity oscillates over certain frequency ranges. In the time domain, the p100 and p300 component of the occipital-parietal ERP are shown to be modulated by the skill level of the population. In the sensor space, a significant difference in p100 amplitude was found with skilled players showing much greater amplitude. However, significant difference in p300 were not found. Multi-variate pattern analysis revealed peak decoding accuracy between skilled and unskilled players occurring over the first 300ms of the response. In the source space, utilizing dSPM in visual cortex labels, both p100 and p300 amplitude differences were found. Increasing the spatial resolution of the activity, enhanced the observable differences between them. With further evidence from MVPA, it appears that a key differentiator between skilled and unskilled players is the early temporal dynamics of the visual system. Specifically, an enhanced early response to the same visual information, displayed by increased p100 and p300 amplitudes in skilled players, facilitating an improved performance. In the frequency domain, there was a difference in pre-stimulus alpha across both populations, with the source localized to occipital regions. Alpha power was significantly higher in unskilled players and showed a strong inverse relationship to reaction time. Theta power during the post stimulus period was significantly higher in unskilled players and also showed a strong inverse relationship to performance, with its source localized to frontal central regions. Low oscillatory power was associated with short reaction time.

Two experimental conditions were used in the Esports aiming test, comparing active and passive conditions on the same AimLabs tasks. In passive conditions, participants would watch a video of a professional player complete the same task before playing it themselves. It served to solidify instructions and display the movement speed required to achieve a high score. In active conditions, players would then take part in the task, implementing what they had just observed. Theta power was significantly higher in active conditions compared to passive conditions in frontal-central sensors and alpha power was higher in occipital-parietal sensors however this effect was only present in skilled players. There were no significant differences found in neural oscillations between populations in the passive condition. Theta power in frontal-central sensors was higher in skilled players during execution of fundamental aiming movements. Through correlational analysis, a positive relationship was discovered between theta power and performance, with high power being associated with high scores in skilled players. Although this relationship was directionally the same in unskilled players, the relationship was not significant but was trending in certain tasks. Alpha power was found to be significantly higher in occipital parietal electrodes in skilled players in the tasks, there were no significant differences in alpha power found in unskilled players. Correlational analysis revealed significant relationships between alpha power and performance with a strong, positive and significant relationship, showing high power and high performance, was found.

### Time domain components and performance

One of the major neural correlates of performance was displayed as p100, the visually evoked potential research, a positive potential detected in occipital sensors 100ms after a visual stimulus onset, as an early electrophysiological correlate of target-orientated visual processing (Desmedt and Robertson 1977). However, by presenting a visual stimulus in different locations in the visual field, significant differences in amplitude and latency in the waveform become apparent (Saba et al., 2023). Increases in target size and a participant’s visual acuity also modulate the amplitude and latency of the p100 waveform, with decreases to latency and gradual increases in amplitude (Li et al., 2011). During continuous visual stimulation, steady state visually evoked potential amplitude increases, but time-locked averages decrease (Rosenstein et al., 1994). Early-stage visual processing also has a key contribution to latter stage, more complex cognition. This occurs in both explicitly and implicitly vision-based behaviours. A notable explicit vision-based behaviour is face perception. This process begins early, reflected in the p100 wave form as a face-selective response (Herrmann et al., 2005). The p100 (referred to as an M100 using MEG) amplitude correlates with successful characterization of face stimuli, discriminating them against other visual stimuli, but not with successful individual face recognition (Liu et al., 2013). Sustained spatial attentional mechanisms are also influenced by visually evoked potentials. By presenting a visual target either to the left or the right visual field immediately preceding a continual visual stimulus (in this case, visual gratings), transient p100 amplitudes were enhanced by sustained attention (Di Russo and Spinelli, 1999). Although sustained attention is modulated by a wide variety of factors, not least neural oscillations, p100s served as an early marker of attention in the time domain.

In the present study, significant increases in p100 amplitude of skilled players might facilitate performance increases in two different ways. Firstly, primary visual detection of the stimuli occurred more strongly in skilled players, denoted by increased amplitude. As such, visual information, even in a basic state is being detected and responded to more strongly in skilled players. Secondly, if p100s contribute to later stage complex cognition, this information is being fed into higher-order areas of the visual cortex to facilitate better target detection. As such, higher p100 amplitude might be driving an enhanced p300 response found in the source space. Peak p300 amplitude was largely similar across both populations in the sensor space but differed in the source space. Although minor differences were found in the sensor space, transformation to the source-space enhanced them whilst also retaining the p100 difference. This suggests that there may be some relationship between p100 amplitude and p300 amplitude whereby together, they impact visual processing and help facilitate a higher level of performance. Therefore, early-stage time-domain voltage dynamics is an important contributor to visuomotor performance and is a substrate through which we can differentiate players based on skill level.

This finding is supported by a multitude of research focusing on p300 which regards this component as a reflection of conscious access (Rutiku et al., 2015)) . Analysis of current spectral density shows that the p300 component, induced by visual stimuli, is highly localized to occipital parietal regions (Ji et al., 1999), supported by the current research. It is induced by visual information since it is abolished by blurring of the visual stimulus (Heinrich et al., 2010) and the source strength during p300ms in occipital parietal sensors is significantly higher during seen trials, as opposed to unseen trials (Babiloni et al., 2006). The present study replicates these findings and proposes an interesting idea for how successful responses are achieved. It is possible that failed trials occur because the visual information was not consciously accessed either at all, or in enough time for the participant to accurately respond. Each trial has a time limit (1000ms) which is incredibly short and a departure from the methodology employed by other visuomotor tasks. This choice implemented an important sporting, but especially Esports, concept of time pressure. Players in sport must react with both speed and accuracy under pressure. As expected, to respond successfully, a participant must consciously access this information quickly. Skilled players can consciously access the salient visual information more frequently, facilitating the improved performance. It is possible that the speed of response is increased by a stronger p100 amplitude, inducing this early-stage visual detection leading to enhanced conscious access, displayed by an increased p300 amplitude. By combining these results, it suggests that skilled players respond faster and with increased accuracy by evoking an enhanced occipital-parietal response over the early stages of visual processing.

However, many of the studies used in support of this research are simple visual detection tasks, asking participants to report if they had seen a stimulus or not. The present task is not so simple. Participants have 1000ms during a trial and it would be unlikely to suggest that a stimulus was not detected at all within that period. Ultimately, there is still a debate in the field about whether p300 reflects conscious awareness or unconscious perception in occipital parietal electrodes, since stimuli are perceived even when observers are unaware of the stimuli (Merikle et al., 2001). P300 theories have been updated to suggest a post-perceptual marker of conscious access not simply conscious perception (Pitts et al., 2014). Therefore, there are still many steps required by the participants to execute a precise movement in time. Training interventions in older populations, have been created displaying enhanced p300 amplitude which was associated with increased cognitive performance (Yang et al., 2018). Skilled players, many of which are experienced in Esports, might display pre-trained enhancements to p300 amplitude which are captured in this study. Whilst this might facilitate an increased visual perception and overall performance, it does not linearly increase reaction time necessarily. The study did not note any differences in the amplitude or latency of p300 between the skilled and unskilled population in the sensor. It is possible to conclude that the visual stimulus more readily enters conscious awareness of skilled players which allows for the utilization of this information to produce a conscious action more frequently. That is, it is not just the amplitude difference of p300, observed in the source-space, that causes the performance difference, but the frequency a high amplitude component is induced which produces the observed performance difference. There could also be further downstream modulations that reflect more complex cognitive/motor, processes and these are only an early indication of conscious perception or awareness. How this information is utilized further is unclear and requires further investigation.

### Neural oscillations and performance

There is a wide array of research isolating the role of alpha power in visual perception research. However, the present study provides evidence for a relationship between pre-stimulus alpha power and complex movement performance. The study found that low pre-stimulus alpha power predicted performance, showing a strong relationship to reaction time. In both populations, there was a significant correlation between low alpha power, and short reaction time.

The study largely supports this theory in several ways. Firstly, skilled players, who more readily execute a successful response, also display a reduction in alpha power which strongly correlates to performance, as has been well established in the research literature. Behaviourally, although reaction time is faster overall in skilled players, this only covers successful trials. The real performance difference is the vastly reduced number of errors in skilled players. As such, the reduction in occipital-parietal, pre-stimulus alpha, which has a significant relationship to reaction time, is present more frequently, leading to a reduction in errors. This inverse relationship between pre-stimulus alpha and performance has been repeatedly found across numerous studies (Van Dijk et al., 2008; Chaumon and Busch, 2014, Benwell et al., 2017; 2022). This decrease in alpha is suggested to facilitate an increase in global excitability (Lange et al., 2013) and reflect the accumulation of evidence over time (Kloosterman et al., 2019) by reducing the threshold required to initiate a decision (Limbach and Corballis, 2016). The present study would support this idea since lower alpha power might release the visual system, direct attention towards the target and induce a motor decision output quicker, facilitating faster reaction time, but also reduced number of errors.

While research has not predominantly used such complex motor outputs, there are several studies which would support this notion. It has been found that pre-stimulus alpha is decreased prior to the execution of successful putts in golf, relative to power in unsuccessful putts (Bablioni et al., 2008) with the strongest alpha power reductions occurring over sensorimotor areas. Golf, however, is a much less visually dominant sport than Esports since players do not adapt to rapidly changing visual stimuli. Although this result slightly differs in location to the current study, with source localization positioning maximal alpha power decreases in the visual cortex, it proposes a link between alpha and performance of a complex movement in sport. Within more visually dominant sports, such as air-pistol shooting, a common feature of high performance is a reduced, pre-stimulus alpha power in occipital regions that precedes successful shots (Kerick et al., 2004; Del Percio et al., 2009). The authors in these cases have postulated decreases alpha power corresponds to an increase in attentional resources used by the visual system assisting performance. The present study strongly supports this idea due to the association between higher performance and low pre-stimulus alpha.

Post-stimulus theta power displayed a strong inverse relationship to performance, in a similar vein to pre-stimulus alpha power. That is, low post-stimulus theta was strongly correlated to high performance. A potential explanation for this general increase is related to realizing the need for cognitive control (Cavanagh and Frank, 2014), coordinated by theta power in the anterior cingulate cortex (ACC) a region commonly linked to theta power (Cohen and Donner, 2013). Its statistical relationship to reaction time has repeatedly been identified (Cohen and Cavanagh, 2011; Cohen and van Gaal, 2014) however these studies often focus on conflict resolution as opposed to more explosive visuomotor performance like the present study. Regardless of trial outcome, theta power increased drastically after stimulus onset across both populations. Interestingly, theta was lower in skilled players. In unskilled players, there was very little difference in peak theta power, but a higher average theta power. Perhaps the sustained increase in unskilled players, and peak difference in skilled players explain a similar phenomenon. That is, moderations to cognitive control were important for high performance. It is possible that unsuccessful trials induced inappropriately large signaling for control that impinged performance in unskilled players. Furthermore, skilled players are more adapted to the movements induced by this task e.g. simple, discrete responses to simple, static stimuli. Perhaps the cognitive demand for this task is not signaled as hard during the post-stimulus period in skilled players as it is in unskilled players, who are potentially naiver to this type of movement. As such, it appears that skilled players regulate the demand for cognitive control where it is appropriate and in a simple, albeit fast, task like this, unskilled players over activate this demand, leading to a reduction in performance.

### Neural oscillations and Esports

Within the Esports aiming task, two different neural correlates became apparent, frontal midline theta power, and occipital-parietal alpha power. Both show a strong positive correlation to performance and an increase in active compared to passive conditions. The increased theta power in both participant populations was observed in the frontal midline of the cortex. This theta pattern is associated with activity in the anterior cingulate cortex (ACC) and prefrontal cortex in source localization studies (Onton et al., 2005; Ishii et al., 2014). Theta power localized to the ACC and other frontal structures has a diverse range of functions in complex cognition such as action regulation (Luu and Pedersen, 2004) and monitoring (Cavanagh et al., 2009); conflict monitoring (Botvinick et al., 2004), task selection (Womelsdorf et al., 2010). Theta power is also highly prevalent in the hippocampus and its thus strongly associated to elements of memory (O’Keefe and Reecee, 1993; Buzsáki, 2005). In particular, retention (Jensen and Tesche, 2002) and encoding (White et al., 2010). However, the present study observes theta power during a highly complex and dynamic visuomotor task. The ACC has strong connections to the motor system (Deiber et al., 1999) and parts of the ACC have been shown to play an essential role in the preparation and readiness (Cunnington et al., 2003), planning (Jankowski et al., 2009) and initiation (Hoffstaedter et al., 2013) of intentional movements. The present study induces a highly complex array of sequential movements requiring precise motor control, speed and accuracy, under time pressure. The extreme temporal density of movements and the precision requirements of them, go far to explain the upregulation of theta power in both populations. As such, it seems that the extreme difference in performance observed between skilled and unskilled players is at least in some part, coordinated by enhanced theta power facilitating increased information flow in the midbrain to the motor cortex, providing the drive for a high, sustained level of performance.

Other studies have linked video-game play to increases in theta power (Pellouchoud et al., 1999) with increases compared to rest periods. The relative power of theta also increases as, time spent playing video games increases or as rounds progress (Sheikholeslami et al., 2007). Theta increases have been shown during violent events in other FPS video games (Salminen and Ravaja, 2008) which required the execution of an aiming movement to hit the opponent. Enhancements to a variety of cognitive abilities of players are commonly associated with video game play including cognitive flexibility (Valls-Serrano et al., 2022), visual processing speed (Kowal et al., 2018) and visual working memory (Seya and Shinoda, 2016). Crucially, top-down attention is also enhanced through video-game play (Chisholm and Kingstone, 2012, However, theta band activity is also linked to cognitive control, signaling an increase in demand (Cavanagh and Frank, 2014). Not only is a high level of attention required, but cognitive and motor control. Both populations are under extreme cognitive load due to the intensity of tasks, speed of movements required, visual complexity and time-pressure.

Importantly, training in older populations, induces significant increases to theta power that are sustained for long periods and are associated with other cognitive test improvements (Anguera et al., 2013). Crucially, the older population of participants were inexperienced with video games and the associated increases in theta were because of video-game play. This highlights the interesting cognitive challenge of Esports since they require significant sensorimotor transformation to the virtual world where movements on the screen mismatch the movements in the real world. For example, moving aim location vertically requires participants to move horizontally in the real world. This is an unusual, non-natural movement and might test the ability of players to adapt. The network required for this type of process is explicitly activated by mouse manipulation (Gorbert et al., 2004), the key movement in Esports aiming. As such, it appears that complex aiming movements, required in Esports, activate a cortical network of prefrontal and premotor cortices, coordinated by theta power. This network operates differently as a function of skill level, with higher skilled, better performing players showing greater activation.

The observed increase in theta power was to be expected and was correctly predicted, especially its strong relationship to performance. However, the relationship of alpha to performance, increasing in active conditions, was not. A possible explanation for this comes from a key finding within cognitive performance tests in video game players. That is, the ability to task switch (Boot et al., 2008), an effect broadly coordinated by attention. Within the neuroscience literature, task switching is commonly associated with a suppression of alpha power and event related desynchronisation in the alpha frequency band prior to movement onset. In the aiming tests, participants must switch between a multitude of competing targets rapidly but also accurately. It might be assumed that, based on the strong field of research on posterior alpha power desynchronisation prior to task switching (Verstraeten and Cluydts, 2002. Sauseng et al., 2006), that these tasks would be associated with decreased alpha power and an inverse relationship to performance would be found. However, the opposite appears to be true. In tasks where there is a high prevalence of switching between tasks, there is a significant, positive relationship to performance.

There are several possible explanations for this result. Firstly, the literature on task switching using event locked methodologies, whereby decreases in posterior alpha are pre-stimulus presentation, which in turn predicts task switching. The present study cannot use these methods and as such cannot make a comment on when alpha power was either high or low, just that over the entirety of the experiment, it remained higher in active conditions and showed a strong relationship to performance across both populations. Also, this switch is voluntary, not determined by a performance variable. In the present study, participants must switch target once it has been ‘destroyed’ either by shooting it or remaining within its boundary for a certain period. As such it is not a voluntary switch but determined by what is required by the task. Furthermore, the tasks that involve switching are associated with additional targets present on screen at any one time, or that are followed incredibly quickly based on performance or time. This is to accurately mimic ecologically valid Esports aiming scenarios where multiple targets are presented at one time, or incredibly close together. As such, the observed power increases in alpha, and their significant positive relationships to performance, might in fact reflect increases in attention to inhibit distractor influences (e.g. alternative targets) and focus attention on the present stimulus. This is well documented in the literature (Benedek et al., 2014). Furthermore, alpha power has been shown to decrease in response to moving images but increase in response to still (Simons et al., 2003). Although this does not fully explain the present data, it offers some explanation for why alpha power increased during multi-target sequential movement tests. Unfortunately, with the methodology employed, limited by the constraints of using a commercial aim-trainer, event related dynamics are not possible to isolate. Nor is any time-frequency decomposition. Future studies should focus on this difference by creating multi-target, complex movement tasks, that incorporate both static and moving targets trials, single and multi-target trials signaled by a trigger system.

## Conclusions, practical implications and future research

The present study utilizes a complex visuomotor performance task to isolate the neural correlates of high performance, examining the differences between a skilled Esports population and an unskilled population. It has uncovered several key contributing factors differentiating between skilled and unskilled players. Across both populations, fast reaction times are correlated to a reduced number of errors. That is, fast reaction times are strongly related to a lower error index, suggesting better performing players are both fast and accurate with their responses. Skilled players display enhanced early-stage visual processing displayed by higher amplitude p100 and p300 components in both the sensor and source space facilitating both enhanced visual perception speed and conscious access to task-related sensory information compared to unskilled players. Furthermore, skilled players display significant reductions in pre-stimulus alpha power. As such, skilled players display a reduction in cortical excitability, coordinated by alpha power in occipital parietal regions, which disengages attention of task-irrelevant information. Unskilled players appear to attend greatly to the fixation cross during the fixation period, resulting in significantly longer saccade latencies in successful trials and prevents fast task switching to the target. This is supported by the early-stage differences in response whereby skilled players display a significantly increased p100 amplitude and decoding scores classifying the response differences, peak over this period in successful trials. Skilled players also display recued post-stimulus theta power, perhaps signifying an increased neural efficiency and ability to manage a high cognitive load.

During the performance of high frequency, complex and sequential Esports-related movements, significant differences occur between skilled and unskilled players. Skilled players show a substantial increase in score across all aiming tasks, and faster TTK. These individual movements are intrinsic to performance in Esports competition are tested here in a controlled and ecologically valid way. AimLabs functions as an aim-training software, commercially available but used by professional players to precisely train aspects of these movements, notably the speed and accuracy. In the present study, the enhanced performance of skilled players appears to be coordinated by frontal theta power emerging from frontal-central regions, presumed to emanate from the anterior cingulate cortex. Theta power, is significantly higher in the active phase of the experiment in skilled players but across both populations, is significantly higher in active versus passive conditions, something consistent with the literature. The higher frontal theta power is associated with increased performance, with stronger associations in skilled players than unskilled players. Furthermore, alpha power in occipital-parietal regions also appears to play a key role, whereby there is a significant positive relationship between alpha power and performance. These relationships are stronger in skilled players. This is presumed to occur by increasing attention on the target being responded to preventing distraction by gating visual encoding of information in peripheral regions. As such, skilled players appear to achieve a higher performance due to an increase in theta power emanating from the ACC which improves the speed and accuracy of motor output and modulate alpha coordinated attention potentially reflecting an increased ability to actively suppress distractors.

Ultimately, the differences between skilled and unskilled players are identifiable using cognitive electrophysiology and Esports functions as a precise, ecologically valid model to use as a sporting framework.

### Applied implications

An important aspect of the present study is relating brain activity to performance using the conceptual framework of sport. As such, understanding how this research could be utilized by both athletes and coaches is important. Crucially, this study is useful not just for Esports, the sport used throughout, it can also act as a model sport with transferrable results to any sport relying heavily on rapid sensory processing and precise movements. Since the study primarily uses Esports as the framework for sporting performance, Esports athletes and coaches will be able to directly use the data to address performance issues. However, this study serves as a blueprint for detecting performance in sensory information driven, fast and precise movements for a wide variety of sports. A key finding is that early-stage visual processing pre and post stimulus onset, directly contributes to the execution of a precise mouse movement. Esports athletes are repeatedly executing this type of movement throughout each game. Yet, this process is implicated in a wide variety of sports too. For example, racket sports such as tennis and badminton, batting sports such as cricket, ball sports such as football and rugby, combat sports such as boxing and MMA. These sports all require very fast visual information processing of target location. It doesn’t matter whether the target is a ball, a limb or an opponent, sports where visual perception is an integral component that drives a motor response will greatly benefit from understanding this research. Therefore, coaches could measure their players brain activity and identify if these components are lacking. It may be that before a stimulus onset, they display high levels of alpha power or, post stimulus onset, they display reduced amplitude of p100 and p300 components. As such, a training intervention could be used to improve these important biomarkers of performance.

In general, there are two predominant ways it can be used for athletes of Esports and other sports, identifying sensory processing weaknesses and training the responses. Identifying weaknesses with the very early phases of visual processing, could be a key indicator that a player has a weakness in this area. This process is integral to many sports and thus the approaches to identify differences across players in an elite squad could benefit them in the future. Although training interventions to modulate time domain components are not well established except through behavioural training, neural oscillations can be modulated through neurofeedback. Through neurofeedback training, it would be possible to reduce alpha oscillations pre stimulus onset. Together with sport-specific coaching, the athlete can implement the feedback training midgame to improve performance. Furthermore, younger players, those on a pathway to becoming professional could be tested as a means of predicting their future potential. We do not know if it is possible to boost early time domain components such as p100/p300 and if that will improve performance directly. Therefore, identifying athletes with these traits that are either innate or, more likely, been developed over their early years, could greatly improve the accuracy of identifying future elite players. The tasks used within this thesis could easily be adapted or changed to include more sport specific information but in their current form could provide benefit to a wide variety of players, boosting their cognitive control, information processing and movement execution.

### Future research

The study utilizes the inherent advantages of using Esports as a model for sport neuroscience in a way that has never previously been achieved. It highlights how Esports and skilled Esports players display a greatly enhanced performance level that is correlated strongly to various aspects of their brain activity. However, further research should focus on neural correlates during the most demanding tasks. The literature is saturated with psychophysics tasks testing a wide variety of cognitive processes. However, the complex nature of Esports, on visual processing and motor responses, is rare. Isolating small elements of cognition might not continue to yield results in the same way as the past. Esports presents a unique opportunity to manipulate complex cognition within the framework of sport, something previously thought impossible. A key advancement would be to isolate evoked brain activity, as opposed to induced (measured in the present thesis). In this way, time-frequency representations of power could be utilized to further isolate how the brains of high skilled players differ from unskilled players. The visual complexity of Esports provides such a novel sensory environment that pushes visual processing to the extreme. Further isolating how this system responds could be a very fruitful avenue.

The data and materials for all experiments are available at: Winstanley, Mike (2024), “ERP and Neural Oscillations: Esports”, Mendeley Data, V1, doi: 10.17632/z7hxyxzgcw.1. The experiments were not preregistered.

